# *Escherichia coli* BarA-UvrY regulates the *pks* island and kills Staphylococci via the genotoxin colibactin

**DOI:** 10.1101/2021.12.17.473262

**Authors:** Jun Jie Wong, Kelvin K.L. Chong, Foo Kiong Ho, Chee Meng Benjamin Ho, Ramesh Neelakandan, Pei Yi Choo, Damien Keogh, John Chen, Chuan Fa Liu, Kimberly A. Kline

## Abstract

Wound infections are often polymicrobial in nature and are associated with poor disease prognoses. *Escherichia coli* and *Staphylococcus aureus* are among the top five most cultured pathogens from wound infections. However, little is known about the polymicrobial interactions between *E. coli* and *S. aureus* during wound infections. In this study, we show that *E. coli* kills *S. aureus* both *in vitro* and in a mouse excisional wound model via the genotoxin, colibactin. We also show that the BarA-UvrY two component system (TCS) is a novel regulator of the *pks* island, which acts through the carbon storage global regulatory (Csr) system. Together, our data demonstrate the role of colibactin in inter-species competition and show that it is regulated by BarA-UvrY TCS, a previously uncharacterized regulator of the *pks* island.

## Introduction

Chronic wound infections are often biofilm-associated and polymicrobial in nature (James, Swogger et al. 2008, Kirketerp-Moller, Jensen et al. 2008, Ammons, Morrissey et al. 2015, Wolcott, Hanson et al. 2016). Polymicrobial wound infections are associated with heightened inflammation and delayed wound healing as compared to monomicrobial wound infections (Kelly 1978, Roy, Elgharably et al. 2014). Within polymicrobial communities, interspecies interactions can increase the pathogenicity of either or both species, inducing virulence gene expression, enhancing growth, or promoting antibiotic tolerance and immune evasion (Weigel, Donlan et al. 2007, Ramsey, Rumbaugh et al. 2011, Pastar, Nusbaum et al. 2013, Tien, Goh et al. 2017). Polymicrobial interactions can also be antagonistic via outcompetition for critical nutrients, by interfering with quorum sensing of competitors (Winter, Thiennimitr et al. 2010, Stacy, McNally et al. 2016, Piewngam, Zheng et al. 2018) or by producing antimicrobial agents to kill competitors (Cotter, Ross et al. 2013, Coulthurst 2019, Sgro, Oka et al. 2019). Antimicrobial agents involved in competitor outcompetition include secreted bacteriocins and effector toxins, which are delivered via specialized secretion systems (Cotter, Ross et al. 2013, Coulthurst 2019, Sgro, Oka et al. 2019).

The most frequently cultured bacterial species from wound infections include *Staphylococcus aureus, Pseudomonas aeruginosa, Enterococcus spp*., *Escherichia coli*, and *Klebsiella pneumoniae* (Citron, Goldstein et al. 2007, Trivedi, Parameswaran et al. 2014). The mechanistic basis of polymicrobial interactions in wounds has been examined for *S. aureus* together with *Pseudomonas aeruginosa* and *Enterococcus faecalis* (Weigel, Donlan et al. 2007, Korgaonkar, Trivedi et al. 2013, Pastar, Nusbaum et al. 2013). However, despite the fact that *E. coli* and *S. aureus* are among the top five most prevalent pathogens in often polymicrobial surgical, diabetic and non-diabetic wound infections (Giacometti, Cirioni et al. 2000, Citron, Goldstein et al. 2007, Trivedi, Parameswaran et al. 2014, Bessa, Fazii et al. 2015), and coexist within diabetic wound microbiomes (Sloan, Turton et al. 2019, Jnana, Muthuraman et al. 2020, Verbanic, Shen et al. 2020), polymicrobial interaction studies between these organisms, or of *E. coli* within wound infections in general, are scarce. We have previously shown that *E. faecalis* promotes *E. coli* biofilm growth and virulence *in vitro* and in a mouse excisional wound infection model (Keogh, Tay et al. 2016). However, the mechanistic basis of interactions between *S. aureus* and *E. coli* remains largely unknown.

In this study, we show that *E. coli* antagonizes *S. aureus* in biofilms and planktonic growth. Both the *E. coli pks* island and the BarA-UvrY two component system (TCS) are required for killing *S. aureus*. The *pks* island encodes the biosynthetic machinery to produce colibactin, a genotoxin that causes DNA damage in eukaryotic cells and is associated with human colorectal cancer (Nougayrede, Homburg et al. 2006, Buc, Dubois et al. 2013). Here we show that *E. coli* colibactin kills *S. aureus* by causing irreparable DNA damage. *E. coli* also antagonizes the growth and survival of *S. aureus* upon co-infection in a mouse excisional wound model, and this antagonism is dependent on the *pks* island and the BarA-UvrY TCS. Finally, we show that the BarA-UvrY TCS regulates the expression of the *pks* island, through the Csr system. Taken together, our data demonstrate the mechanism by which *E. coli* colibactin acts in inter-species competition to kill *S. aureus* during wound infection.

## Results

### *E. coli* antagonizes the growth of *Staphylococcus* species during *in vitro* co-culture

To investigate the mechanistic basis of interactions between *S. aureus* and *E. coli*, we assessed the growth of each species within macrocolony biofilms and planktonic co-culture, followed by enumeration of viable CFU of each species on selective media. We first grew dual species macrocolonies with *E. coli* UTI89 partnered with one of six different strains of *S. aureus*. By 24 hours, the CFU of all *S. aureus* strains tested, including the methicillin resistant *S. aureus* strain (MRSA) USA300, fell from the initial inoculum of 1×10^5^ CFU to below limit of detection in the presence of *E. coli*, demonstrating that *E. coli* UTI89 can kill *S. aureus* within macrocolony co-culture (**Figure 1A**). Conversely, not all *E. coli* strains tested could kill *S. aureus* strain HG001; *E. coli* MG1655 did not kill *S. aureus*, suggesting that this phenotype is specific to certain *E. coli* strains (**Figure 1B**). To test whether *E. coli* could similarly kill other members of the Staphylococcus genus, we grew macrocolonies of *E. coli* with *S. saprophyticus* and *S. epidermidis* and observed that both *Staphylococcus* species were killed (**Figure 1C**). We also observed *S. aureus* killing in planktonic culture with *E. coli*; however, the killing was less efficient and took 48 hours to approach the limit of detection (**Figure 1D**). Similar to the macrocolony assay, *E. coli* MG1655 was also unable to kill *S. aureus* during planktonic growth (**Figure 1D**). *E. coli* CFU remained unchanged when grown alone or co-cultured with different strains of *S. aureus* (**Supplementary Figure 1**). These results show that *E. coli* can kill *S. aureus* both within macrocolony biofilms and planktonic cultures.

**Figure 1.**
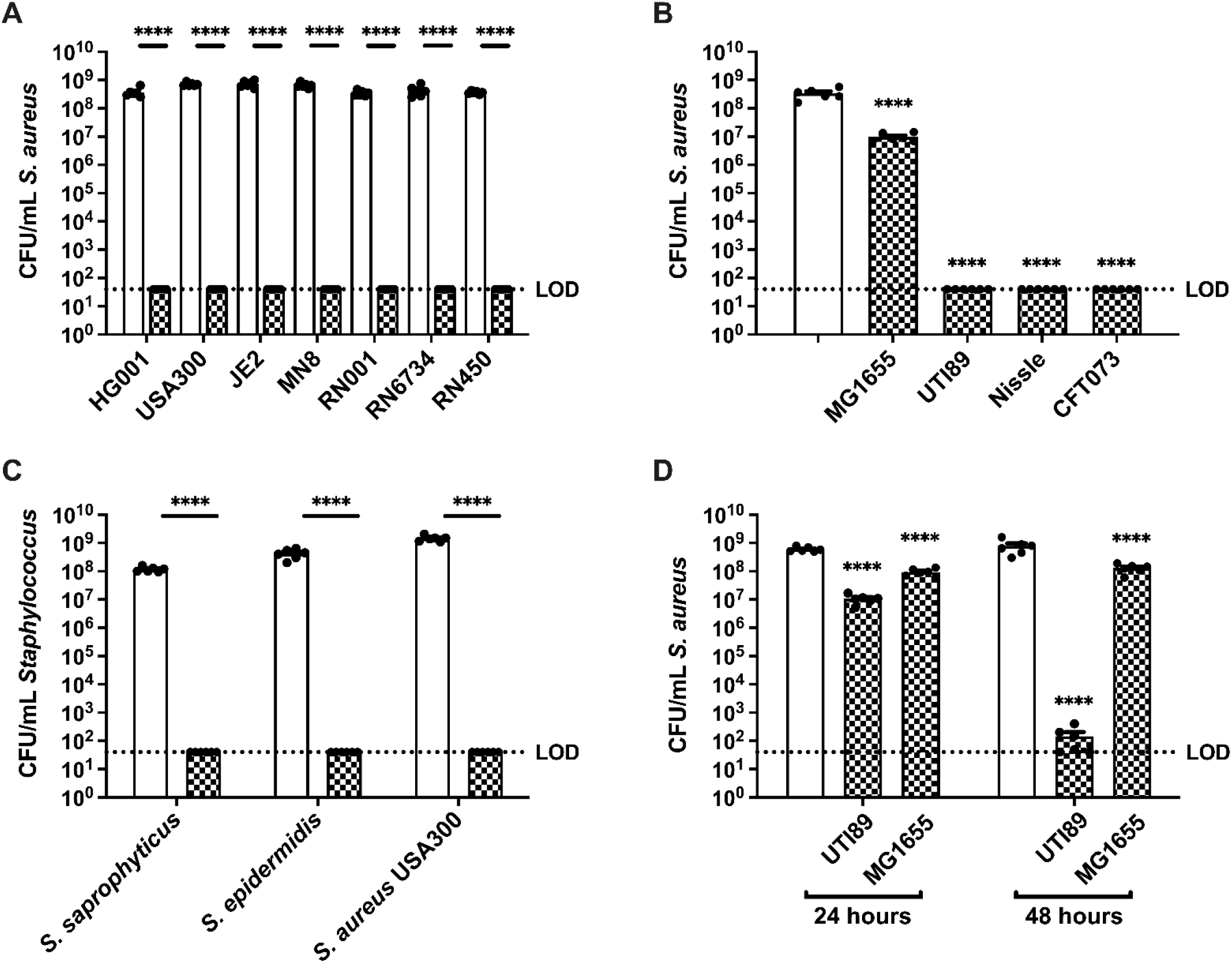
*E. coli* kills *Staphylococcus spp in vitro*. **(A)** Enumeration of different strains of *S. aureus* grown alone or together with *E. coli* UTI89 in macrocolonies for 24 h. N=6 independent experiments. **(B)** Enumeration of *S. aureus* HG001 from single species or mixed species macrocolonies containing the indicated strain of *E. coli*, at 24 h. N=6 independent experiments. Statistical significance was determined by one-way ANOVA with Dunnett’s test for multiple comparison. **(C)** *Staphylococci* from single species or mixed species macrocolonies co-cultured with *E. coli* UTI89 for 24 h. N=6 independent biological experiments. **(D)** Enumeration of *S. aureus* after planktonic growth alone or mixed with either *E. coli* UTI89 or MG1655 for 24 or 48 hours. N=6 independent experiments (**A-D**) Data from single species macrocolonies or planktonic culture are indicated with open bars, and data from mixed species (all inoculated at a ratio of 1_EC_:1_SA_) macrocolonies are indicated with checked bars. Individual data points from each biological replicate are indicated with closed circles. (**A, C, D**) Statistical significance was determined by two-way ANOVA with Sidak’s (A and C) and Tukey’s (D) test for multiple comparisons. ****p< 0.0001. Error bars represent SE from the mean. All statistical tests were performed on log-transformed CFU data. (See **Supplementary Figure 1** for paired *E. coli* CFU to match the *S. aureus* data shown here.)

### *E. coli* does not kill *S. aureus* by prophage induction

To gain insight into the mechanism underlying *E. coli*-mediated killing of *S. aureus*, we examined the gene expression profiles of both *E. coli* and *S. aureus*, comparing single and mixed species macrocolonies. We extracted RNA from macrocolonies grown for 6 hours since viable *S. aureus* could be recovered at the timepoint (data not shown). Several gene expression pattern changes were distinctive **(Figure 2)**. First, genes related to iron acquisition and utilization were induced in both species within mixed species macrocolonies compared to single species macrocolonies, suggesting that iron is limiting during co-culture. Second, genes associated with phage or mobile elements comprised the most highly induced functional category for *S. aureus* within mixed species macrocolonies. In *S. aureus*, the DNA damage-induced SOS response can induce resident prophages, leading to *S. aureus* lysis (Selva, Viana et al. 2009). Since we also observed that genes involved in DNA repair, such as *recA* and *uvrY*, were upregulated in *S. aureus* mixed species macrocolonies, we hypothesized that prophage induction contributes to *E. coli*-mediated killing of *S. aureus*. However, when we tested the prophage-cured *S. aureus* strain RN450 in the mixed species macrocolony assay (Novick 1967), we observed that it was also readily killed by *E. coli* (**Figure 1A**), indicating that prophage induction was not the only mechanism by which *E. coli*-induced death in *S. aureus* occurred.

**Figure 2.**
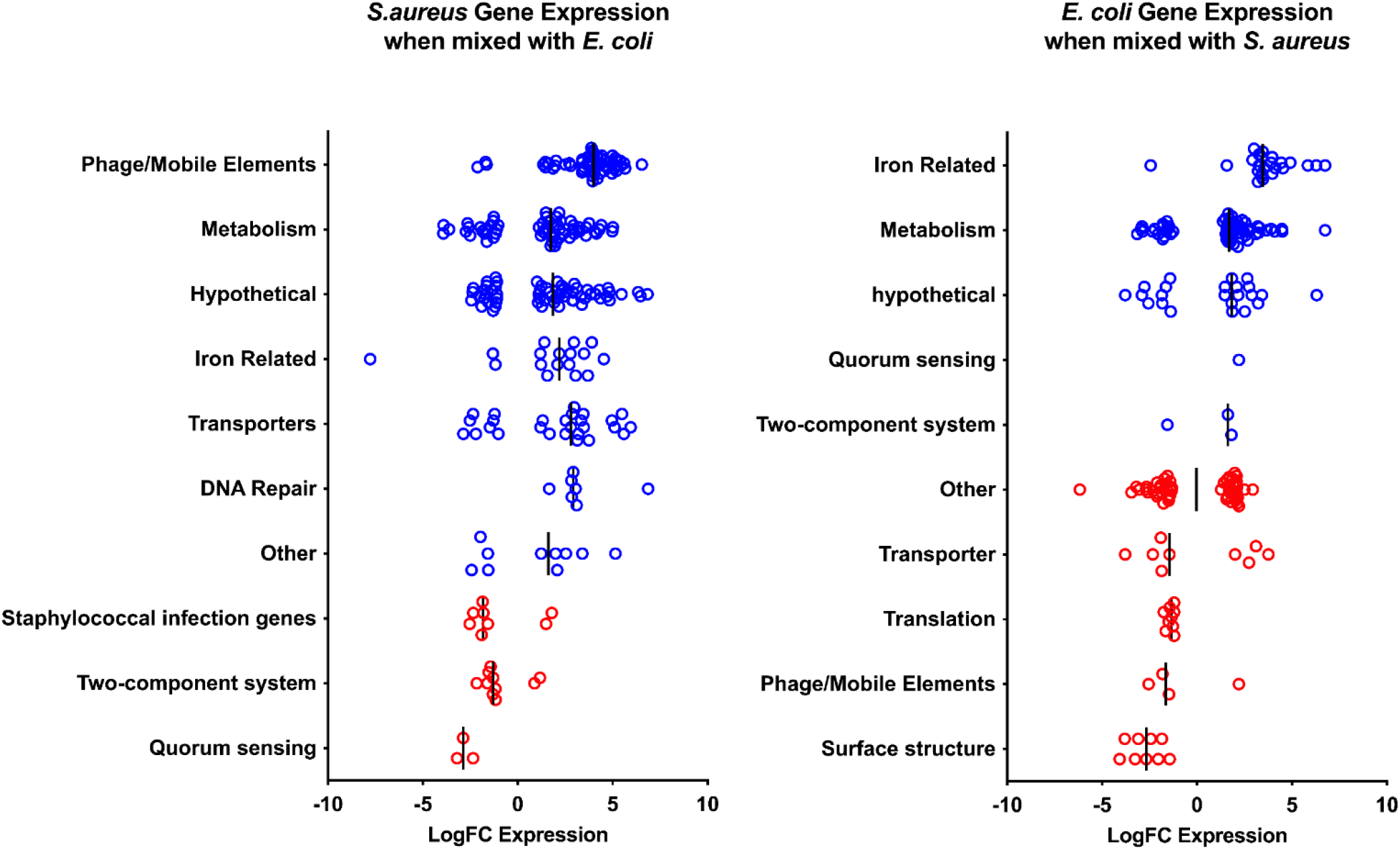
Co-culture of *E. coli* and *S. aureus* promotes differential gene expression. Transcription comparison between single species *S. aureus* strain HG001 or *E. coli* strain UTI89 macrocolonies and mixed species macrocolonies (1EC:1SA) after 6 h incubation, a time point at which we do not observe significant *S. aureus* killing. Vertical black lines represent median values for each gene category. Each circle represents a gene that is differentially regulated (p<0.05, FDR<0.05) in the mixed species macrocolony compared to the single species macrocolony in the respective functional categories, with blue color indicating a functional category where the median value shows increased expression in the mixed species macrocolony and red color indicating decreased expression. Data represent ≥2 biological replicates.

### The colibactin *pks* island and BarA-UvrY TCS contributes to *E. coli* mediated growth antagonism of *S. aureus*

To identify the genes involved in *E. coli*-mediated killing of S. *aureus*, we screened 14,828 *E. coli* UTI89 transposon mutants for failure to antagonize *S. aureus* growth in a macrocolony assay. We validated each *E. coli* gene identified in the transposon screen for its inability to kill *S. aureus* after 24 hours of macrocolony co-culture and the transposon insertion sites for validated mutants were determined by whole genome sequencing. The majority (99 out of 108) of transposon insertions mapped to genes of the *pks* island and genes encoding the two component system (TCS) BarA-UvrY (**Table S1**). To confirm that these *E. coli* loci impacted *S. aureus* survival, we generated deletion mutants comprising the entire *pks* island, as well as for *barA* and *uvrY*, all of which had significantly higher *S. aureus* CFU in the mixed species macrocolony compared to *E. coli* UTI89 wild type (**Figure 3A**). These mutants grew as well as wild type (data not shown), showing that attenuation of the killing phenotype was not due to growth differences. Although co-culture with *E. coli* Δ*pks* did not restore *S. aureus* growth to single species growth levels, *S. aureus* CFU were similar to that in co-culture with *E. coli* MG1655 (**Figure 1A**), which also failed to kill *S. aureus*. We therefore surmise that nutrient competition within mixed species macrocolonies results in a 1-2 log decrease in *S. aureus* compared to *S. aureus* grown alone. Notably, co-culture with either Δ*barA* or Δ*uvrY* only partially restored *S. aureus* growth, suggesting that the growth antagonism is not completely abolished when the BarA-UvrY TCS is inactivated (**Figure 3A**). These data show that the *pks* island is necessary for *S. aureus* killing and suggest that the BarA-UvrY TCS may either directly or indirectly regulate the expression of the *pks* island or the activity of its gene products.

**Figure 3.**
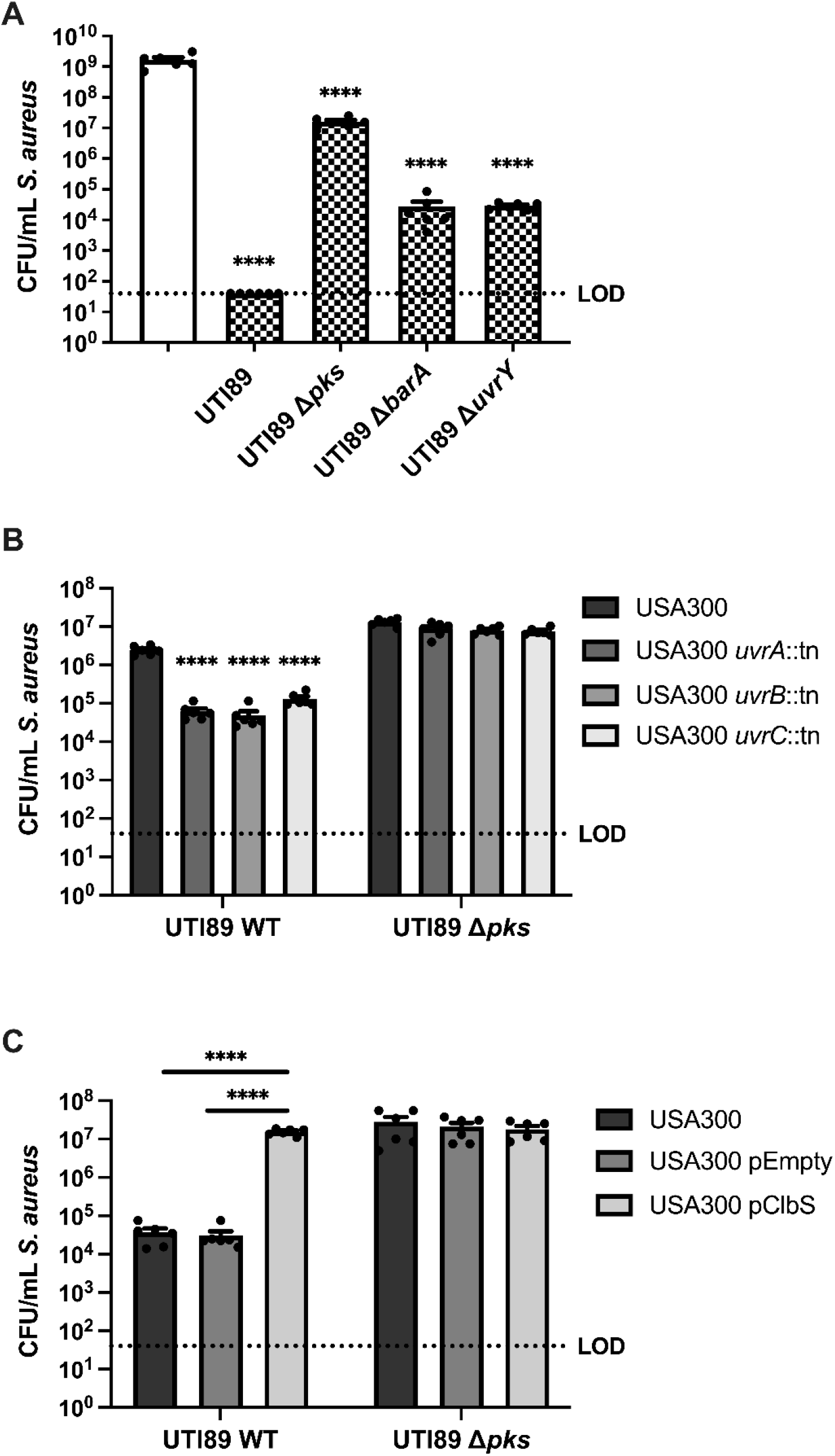
The BarA-UvrY two component system (TCS) and the *pks* island are required for *E. coli*-mediated killing of *S. aureus*. **(A)** Enumeration of *S. aureus* USA300 LAC and mixed (1_EC_:1_SA_) macrocolonies with either UTI89 wild type or knockout mutants of the *pks* island, *barA* or *uvrY*. Data from single species macrocolonies are indicated with open bars, and data from mixed species macrocolonies are indicated with checked bars. N=6 independent biological experiments. Statistical significance was determined by one-way ANOVA with Dunnett’s test for multiple comparison. **(B)** Enumeration of *S. aureus* USA300 LAC from 8 h macrocolonies. Wild type *S. aureus* USA300 LAC and *uvrABC* transposon mutants were mixed 1_EC_:1_SA_ with either *E. coli* UTI89 or knockout mutants of the *pks* island. N=6 independent experiments. **(C)** Enumeration of *S. aureus* from 24 h macrocolonies. Wild type *S. aureus* USA300 LAC was transformed with pJC-2343 (pEmpty) or pJC-2343-ClbS (pClbS) and mixed 1:1 with either *E. coli* UTI89 or knockout mutants of the *pks* island. N=6 independent experiments. Individual data points are indicated with closed circles. **(B and C**) Statistical significance was determined by Two-way ANOVA with Tukey’s test for multiple comparisons. ****p< 0.0001, error bars represent SE from the mean. All statistical tests were performed on log-transformed CFU data. (See **Supplementary Table S1** for full list of transposon mutants identified in this screen.)

### The genotoxin colibactin kills *S. aureus* by inducing DNA damage

The *pks* island encodes enzymes required for the synthesis of the genotoxin colibactin (Fais, Delmas et al. 2018). *E. coli* strains carrying the 54 kb *pks* island generate DNA adducts and induce DNA crosslinks in mammalian cells (Nougayrede, Homburg et al. 2006, Vizcaino and Crawford 2015, Bossuet-Greif, Vignard et al. 2018, Wilson, Jiang et al. 2019). In bacteria, DNA adducts and crosslinks can be repaired via the nucleotide excision repair (NER) pathway, facilitated by the UvrABC endonuclease complex (Kisker, Kuper et al. 2013). Accordingly, *pks+ E. coli* strains lacking both UvrB and the ClbS colibactin immunity protein, which protects *E. coli* from colibactin-mediated autotoxicity, are severely impaired for growth (Bossuet-Greif, Dubois et al. 2016). Consistent with DNA damage, we found that *S. aureus uvrABC* genes were significantly upregulated in mixed species macrocolonies with *E. coli* (**Figure 2**). Macrocolony co-culture of *S. aureus uvrA, uvrB* and *uvrC* transposon mutants with *E. coli* resulted in accelerated *pks*-dependent killing and significantly fewer *S. aureus* CFU at 8 hours, suggesting a role for NER in the protection of *S. aureus* from colibactin-mediated killing **(Figure 3B)**. To further investigate the role of colibactin in *S. aureus* killing, we expressed the colibactin immunity protein ClbS in *S. aureus*. Expression of ClbS in *S. aureus* cells conferred full protection from *E. coli pks*-mediated killing, suggesting that colibactin is responsible for *S. aureus* cytotoxicity **(Figure 3C)**. Collectively, these data establish that colibactin, synthesized by *pks*+ *E. coli*, kills *S. aureus* by causing DNA damage in *S. aureus* in a manner similar to that documented in eukaryotic cells.

### N-myristoyl-D-Asn causes pore formation in S. aureus

Maturation of colibactin requires the removal of the prodrug motif, *N*-myristoyl-D-Asn (NMDA), by ClbP peptidase (Bian, Fu et al. 2013, Brotherton and Balskus 2013). NMDA is the most abundant of the *pks* island metabolites along with its analogues that vary in acyl chain lengths (C_12_ to C_16_) (Vizcaino, Engel et al. 2014). These intermediates do not exhibit cytotoxic or genotoxic activity in HeLa cells; however, NMDA can modestly inhibit *Bacillus subtilis* growth (Vizcaino, Engel et al. 2014). Thus, we investigated if the production of NMDA could provide an alternative explanation for the killing of *S. aureus* by *pks*+ *E. coli*. We synthesized NMDA and added it at increasing concentrations to *S. aureus* but we observed minimal dose-dependent growth inhibition, and only at high concentrations of 600 µM (**Supplementary Figure 2A**). Despite the absence of significant toxicity, NMDA treatment resulted in increased *S. aureus* membrane permeability as measured by propidium iodide uptake using flow cytometry (**Supplementary Figure 2B**). Thus, NMDA, which is the most abundant colibactin metabolite isolated from culture supernatants (Vizcaino, Engel et al. 2014) and is therefore likely released from *E. coli* along with colibactin, can compromise *S. aureus* membrane integrity.

### BarA-UvrY TCS regulates *pks* island genes via the Csr system

Since both *pks* and *barA/uvrY E. coli* mutants failed to kill *S. aureus*, we hypothesized that the BarA-UvrY TCS is involved in the regulation of the *pks* island. To investigate this, we first compared the expression of *pks* island genes (*clbA* and *clbB*) between single species macrocolonies of wild type *E. coli* and *E. coli* Δ*uvrY* and found out that the expression of both *clbA* and *clbB* were significantly lower in the *E. coli* Δ*uvrY* macrocolony (**Figure 4A**). Next, we examined the expression of *pks* island genes in the presence or absence of *S. aureus*, and found that *E. coli clbA* expression was significantly increased in mixed species macrocolonies compared to *E. coli* single species macrocolonies (**Figure 4B)**. By contrast, expression of both *clbA* and *clbB* were significantly lower when *E. coli* Δ*uvrY* was co-cultured in macrocolonies with *S. aureus*, compared to wild type *E. coli*-*S. aureus* macrocolonies, suggesting that the BarA-UvrY TCS is involved in regulating *pks* island gene expression (**Figure 4C**). Moreover, *S. aureus*-dependent induction of *clbA* and *clbB* gene expression in wild type *E. coli* was significantly attenuated upon macrocolony co-culture with *E. coli* Δ*uvrY* (**Figure 4D**), together suggesting that *E. coli* can use the BarA-UvrY system to sense *S. aureus* and induce *pks* island gene expression.

**Figure 4.**
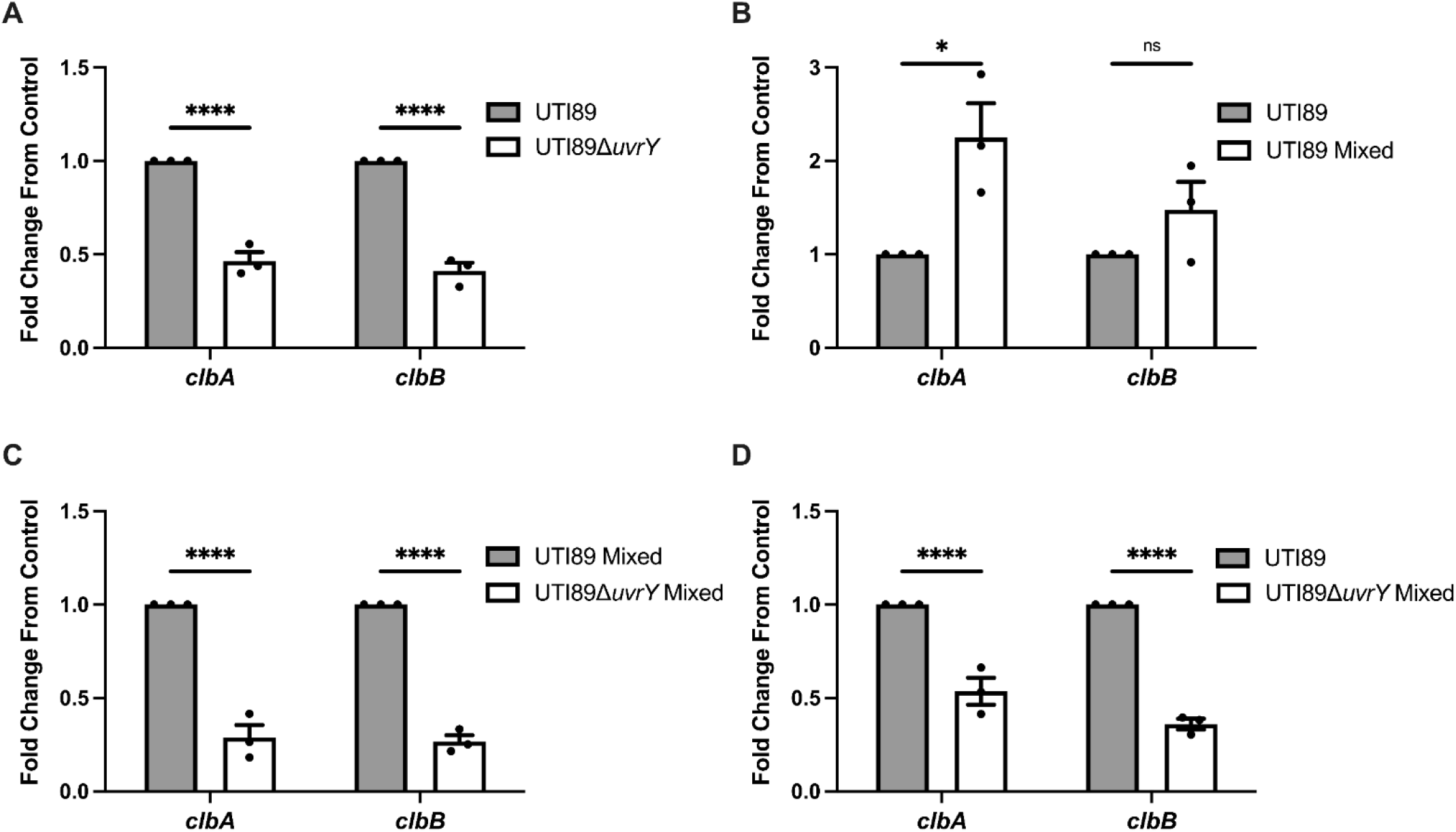
Co-culture of *E. coli* and *S. aureus* induces *pks* island expression in a BarA-UvrY TCS dependent manner. **(A)** RT-qPCR of *E. coli* single species macrocolonies and *E. coli* Δ*uvrY* single species macrocolonies at 24 h. **(B)** RT-qPCR of *E. coli* single species macrocolonies and *E. coli* mixed species macrocolonies at 24 h. **(C)** RT-qPCR of *E. coli* mixed species macrocolonies and *E. coli* Δ*uvrY* mixed species macrocolonies at 24 h. **(D)** RT-qPCR of *E. coli* single species macrocolonies and *E. coli* Δ*uvrY* mixed species macrocolonies at 24 h. N=3 independent experiments, each the average of 4 technical replicates. Gene expression was normalized to the *gyrA* housekeeping gene. Individual data points from each biological replicate are indicated with closed circles. Statistical significance was determined by Bonferroni’s multiple comparisons test for two-way ANOVA, ****p< 0.0001, error bars represent SE from the mean.

The BarA-UvrY TCS regulates the expression of the Csr system, which in turn regulates a variety of metabolic and virulence genes via the global regulator CsrA (Romeo and Babitzke 2018). CsrA is a post-transcriptional regulator that can either promote or suppress gene expression (Timmermans and Van Melderen 2010). Activation of the BarA-UvrY TCS, leads to the expression of the sRNAs CsrB and CsrC, which bind to CsrA and inhibit the regulatory activity of CsrA (Timmermans and Van Melderen 2010). Therefore, if BarA-UvrY regulates *pks* transcription via CsrA, we hypothesized that increasing expression of CsrA would suppress *pks* island gene expression and increasing expression of CsrB would lead to the upregulation of *pks* island genes. Consistent with these predictions, we observed that overexpression of CsrA leads to the downregulation of *pks* island genes, suggesting that CsrA is a negative regulator of the *pks* island (**Figure 5A**). Conversely, overexpression of CsrB resulted in upregulation of *pks* island genes (**Figure 5B**). Co-culturing the *E. coli* overexpression strains with *S. aureus* in macrocolonies was consistent with the *pks* expression data, such that CsrA overexpression led to reduced *pks* gene expression and *S. aureus* killing, and CsrB overexpression led to increased *pks* gene expression and enhanced killing (**Figure 5C**). CsrA regulates gene expression by binding to its target mRNA at GGA motifs, which can be found in the 5’ untranslated region, early coding region, and the stem-loop structure of the mRNA (Dubey, Baker et al. 2005, Vakulskas, Potts et al. 2015, Potts, Vakulskas et al. 2017). We generated a *E. coli* strain where the GGA motifs in *clbR* were modified, which is predicted to reduce CsrA binding efficiency to *clbR* mRNA, as has been reported for other CsrA mRNA substrates (Dubey, Baker et al. 2005, Romeo and Babitzke 2018). We hypothesized that this strain, *E. coli clbR*^mut^ would significantly increase *S. aureus* killing because CsrA suppression of *clbR* expression would be alleviated. Consistent with this hypothesis, we observed that *E. coli clbR*^mut^ kills *S. aureus* faster than wild type *E. coli* (**Figure 5D**). Collectively, these data demonstrate that the *E. coli* BarA-UvrY TCS senses *S. aureus* and responds by inducing *pks* gene expression via CsrA which acts as a negative regulator of the *pks* island.

**Figure 5.**
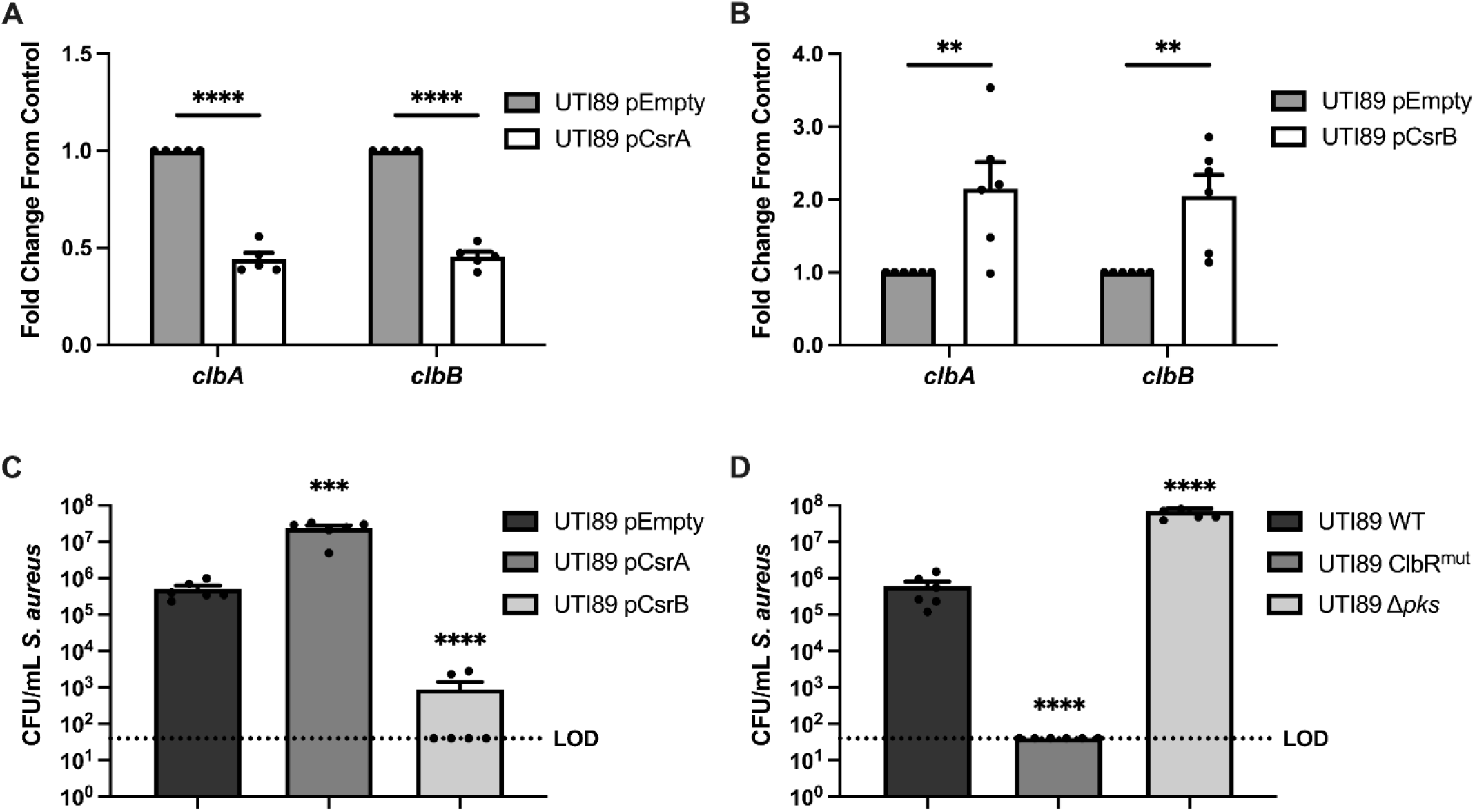
The BarA-UvrY TCS regulates the *pks* island via the Csr system. **(A)** RT-qPCR of 16 h macrocolonies of *E. coli* pTrc99a (pEmpty) and *E. coli* pTrc99a-CsrA (pCsrA). **(B)** RT-qPCR of 16 h macrocolonies of *E. coli* pEmpty and *E. coli* pCsrB. N=5-6 independent experiments, each the average of 2 technical replicates. Gene expression was normalized to the *gyrA* housekeeping gene. Statistical significance was determined by two-way ANOVA with Bonferroni’s test for multiple comparison, **p< 0.01, ****p< 0.0001, error bars represent SE from the mean. ****p< 0.0001, error bars represent SD from the mean. **(C)** Enumeration of *S. aureus* from 16 h mixed macrocolonies (1_EC_:1_SA_) with either *E. coli* pEmpty, *E. coli* pCsrA or *E. coli* pCsrB. N=6 independent experiments. (**D**) Enumeration of *S. aureus* mixed macrocolonies (1_EC_:1_SA_) from 16 h with either *E. coli* WT, *E. coli* ClbR^mut^ or *E. coli pks* deletion mutant. N=6 independent experiments. Individual data points are indicated with closed circles. (**C and D**) Statistical significance was determined by Ordinary One-way ANOVA with Dunnett’s test for multiple comparison. ****p< 0.0001, error bars represent SE from the mean. Statistical tests were performed on log-transformed data.

### *E. coli* antagonizes the growth of *S. aureus* in a mouse model of wound infection

To determine whether *E. coli* could similarly antagonize *S. aureus* growth *in vivo* within a mixed species wound infection, we infected excisional wounds of C57BL/6 mice with 10^6^ CFU each of *E. coli* and *S. aureus* cells and monitored the bacterial burden at the wound site. At 24 hours post infection (hpi), *S. aureus* CFU were significantly reduced when co-infected with *E. coli* as compared to single species *S. aureus* infection (**Figure 6A**). Upon co-infection with *S. aureus* and *E. coli* Δ*pks, S. aureus* CFU remained similar to *S. aureus* single species infected wounds (**Figure 6B**), whereas co-infection *E. coli* Δ*barA* or *E. coli* Δ*uvrY* resulted in increased *S. aureus* survival, but not restoration to single species levels (**Figure 6C**), similar to our *in vitro* results (**Figure 3A**). Together, these data demonstrate that both the *E. coli pks* island and the BarA-UvrY TCS are important for the growth antagonism of *S. aureus* observed in mixed species infections *in vivo*.

**Figure 6.**
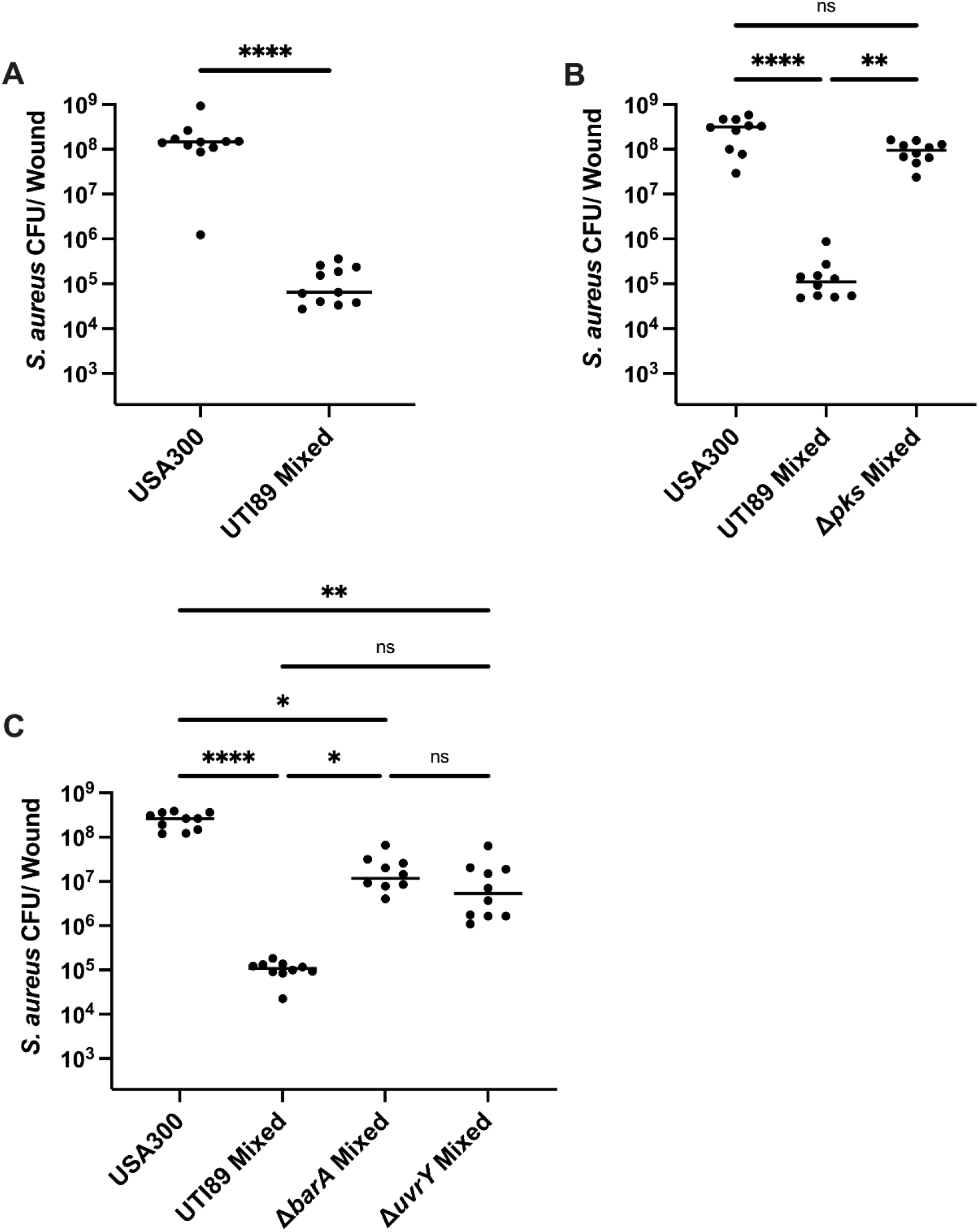
*E. coli* antagonizes *S. aureus* growth during wound infection and antagonism is dependent on the *pks* island and the BarA-UvrY TCS. Mice were co-infected with *E. coli* UTI89 and *S. aureus* USA300 LAC at 1-2 × 10^6^ CFU/wound. Wound CFU were enumerated at 24 h post infection. *S. aureus* single species infection or co-infection with **(A)** *E. coli* UTI89 WT, **(B)** *E. coli pks* mutant, or **(C)** BarA-UvrY TCS mutants. Each black circle represents one mouse, horizontal lines represent the median. N=2 independent experiments, each with 5-6 mice per group. Statistical analysis was performed using Kruskal-Wallis test with Dunn’s post-test to correct for multiple comparisons. *p< 0.05, **p< 0.01, ****p< 0.0001.

## Discussion

*Escherichia coli* and *Staphylococcus aureus* are both important pathogens that cause wound infections, blood infections, urinary tract infections and infective endocarditis (Kaper 2005, Lauridsen, Arpi et al. 2011, Tong, Davis et al. 2015). Both *E. coli* and *S. aureus* can exhibit polymicrobial synergy with other bacterial species during infection, which is advantageous for these pathogens, but often leads to adverse disease outcomes (Pastar, Nusbaum et al. 2013, Keogh, Tay et al. 2016, Tien, Goh et al. 2017). While many studies have investigated the mechanistic basis of polymicrobial interactions between different microbial species, the molecular interactions between *E. coli* and *S. aureus* have not been reported. In this study, we found that *E. coli* production of colibactin is responsible for the growth antagonism toward *S. aureus*, resulting in significant inhibition of *S. aureus in vitro* and *in vivo* during polymicrobial wound infection. *E. coli pks* genes are upregulated during co-culture with *S. aureus*, supporting the proposed role of *E. coli* colibactin as an effector for niche adaptation or domination (Tronnet, Floch et al. 2020). Finally, we found that the *E. coli* two component signal transduction system BarA-UvrY senses the polymicrobial environment leading to the upregulation of the *pks* island via the CsrA system.

Colibactin, encoded by the *pks* genomic island, is a genotoxin that causes double-stranded DNA breaks, chromosomal instability, and cell cycle arrest in eukaryotic cells (Nougayrede, Homburg et al. 2006, Cuevas-Ramos, Petit et al. 2010). A functional role for the *pks* island in polymicrobial interactions has also been reported. Colibactin altered the gut microbiome composition in newborn mice when the pregnant mothers were previously colonized with *pks*^+^ *E. coli*, with Firmicute reduction observed starting 35 days after birth (Tronnet, Floch et al. 2020). More relevant to our work, *E. coli* episomally expressing the *pks* island spotted onto lawns of *S. aureus* gave rise to small zones of inhibition around the *pks*^+^ *E. coli* colonies, although they did not confirm that colibactin was the factor producing antibiotic activity (Fais, Cougnoux et al. 2016). We similarly observed that this antagonism was more efficient within macrocolonies, where *E. coli* completely inhibits the growth of *S. aureus* within 24 hours as compared to planktonic cultures, where we only saw significant growth inhibition at 48 hours. These data indicate that colibactin-mediated growth inhibition of *S. aureus* is favored at close proximity but is not a biofilm dependent phenotype.

Colibactin is genotoxic toward eukaryotic cells, and the ClbS immunity protein, which hydrolyzes colibactin into a non-toxic compound, protects the mammalian host DNA from damage (Bossuet-Greif, Dubois et al. 2016, Tripathi, Shine et al. 2017). Consistent with the role of ClbS as a colibactin immunity protein, *E. coli clbS* mutants displayed increased *recA* expression, and *clbS uvrB* double mutants are significantly attenuated in growth (Bossuet-Greif, Dubois et al. 2016). Similarly, we observed transcriptional induction of genes involved in DNA repair, such as *recA* and *uvrA* in *S. aureus* upon co-culture with *E. coli*. Furthermore, *S. aureus* mutants of the nucleotide excision repair (NER) pathway (*uvrA, uvrB* and *uvrC*) were more susceptible to *E. coli* mediated growth inhibition. Importantly, *clbS* expression in *S. aureus* conferred full protection from *E. coli*-mediated growth inhibition. Therefore, this experiment confirms that colibactin is the factor inhibiting the growth of *S. aureus*, a conclusion that is also consistent with other observations made in this study. First, colibactin is highly unstable (Fais, Delmas et al. 2018), which explains why the most efficient growth inhibition of *S. aureus* is seen when both species are in close contact within a macrocolony biofilm, whereas killing was less efficient in planktonic co-culture. Second, only *pks*^+^ *E. coli* can inhibit the growth of *S. aureus*, which explains why *E. coli* MG1655 is unable to inhibit *S. aureus* growth, as it does not possess the *pks* island (Bonnet, Buc et al. 2014, Yang and Jobin 2014). Finally, while NMDA impaired the integrity of the *S. aureus* membrane and has been shown to modestly inhibit *B. subtilis* growth (Vizcaino, Engel et al. 2014), full protection by ClbS expression in *S. aureus* indicates that mature colibactin, and not an intermediate such as NMDA, is responsible for *S. aureus* killing by *E. coli*. To date, it is not known how colibactin enters mammalian or bacterial target cells; however, the ability of NMDA to compromise the membranes of *S. aureus* could serve as a mechanism for colibactin entry in some circumstances.

In *E. coli, pks* island genes are upregulated when iron is limited in a Fur-dependent manner, while *pks* island genes are downregulated when iron is in abundance (Tronnet, Garcie et al. 2016, Tronnet, Garcie et al. 2017). While gene expression profiling indicates that both *E. coli* and *S. aureus* are experiencing iron-limitation in mixed species macrocolonies, iron-supplementation experiments did not prevent *E. coli*-mediated killing of *S. aureus* (data not shown), suggesting that iron restriction is not the sole driver of *pks* expression in this mixed species interaction. Indeed, we have shown that the BarA-UvrY TCS also regulates the expression of the *pks* island. The presence of *S. aureus* leads to transcriptional upregulation of *pks* island, in a BarA-UvrY-dependent manner. However, while *S. aureus* growth is partially restored when *S. aureus* is co-cultured with either *barA* or *uvrY* mutants, it is not restored to the level of the Δ*pks* strain. These data may suggest that the BarA-UvrY is also not the sole driver of *pks* expression and that, perhaps, iron limitation could impact *pks* island transcription in the absence of the BarA-UvrY TCS.

One of the direct targets of the BarA-UvrY TCS is the Csr system, which in turn regulates diverse functional pathways such as glycolysis, gluconeogenesis, and expression of virulence factors such as biofilm formation, toxin production and pilus expression (Timmermans and Van Melderen 2010). Upon activation of BarA, activated UvrY induces the expression of *csrB* and *csrC* (Suzuki, Wang et al. 2002, Weilbacher, Suzuki et al. 2003). *csrB* and *csrC* are small non-coding RNAs (sRNAs) that bind to CsrA to negatively regulate the activity of the CsrA transcriptional regulator (Timmermans and Van Melderen 2010). The BarA-UvrY TCS have been previously demonstrated to be essential for *pks*+ *E. coli*-mediated genotoxicity of HeLa cells (Homburg 2007), together suggesting a link between this TCS, the Csr system, and colibactin synthesis, which our data support. Colibactin function has largely been studied in the context of colorectal cancer (Fais, Delmas et al. 2018). Since we know that *pks* is regulated by both iron and the BarA-UvrY system, and since iron may not always be a limited nutrient in the gastrointestinal (GI) tract (Seyoum, Baye et al. 2021), these facts suggest that BarA-UvrY may be the predominant regulator of *pks* island expression in the GI tract. Short -chain fatty acids (SCFAs) such as acetate, propionate and butyrate are abundant in the GI tract (Silva, Bernardi et al. 2020). Furthermore, these SCFAs have been demonstrated to be the stimulus of the BarA histidine kinase (Chavez, Alvarez et al. 2010).

As such, the presence of SCFA could serve as a signal for upregulation of the *pks* island via the BarA-UvrY TCS in the GI tract. Consistent with a role for SCFA sensing by BarA, both *E. coli* and *S. aureus* have been reported to accumulate and increase production of these molecules when is iron is limited, which is suggested by gene expression profiles of mixed species macrocolonies (Friedman, Stauff et al. 2006, Folsom, Parker et al. 2014). Overall, these results not only bridge the knowledge gap in understanding the polymicrobial interactions between *E. coli* and *S. aureus*, but also contribute to the understanding *pks* island regulation in the context of polymicrobial infections.

*E. coli* and *S. aureus* are causative agents of wound infections (Negut, Grumezescu et al. 2018), where they can be co-isolated from polymicrobial wound infections (Bessa, Fazii et al. 2015, Krumkamp, Oppong et al. 2020). However, the frequency of co-infection with these two species is not clear from epidemiology literature. Our report that *pks*+ strains of *E. coli* inhibit *S. aureus* growth in wound infections suggest several possibilities. It is possible that when *E. coli* and *S. aureus* are co-isolated from wounds, the *E. coli* strains do not encode or express the *pks* island. Alternatively, *pks*+ *E. coli* that are co-isolated with *S. aureus* from wound infections may retain spatial segregation within wound biofilms such that colibactin is not in close enough proximity to *S. aureus* to severely limit its growth. It is also possible that host-dependent factors may serve to inactivate colibactin in some individuals. More detailed epidemiological, metagenomic, and pangenomic studies of wound infection microbiota are required to understand the ecological landscape within wound infections.

In summary, in this study, we have shown that co-infection with *S. aureus* induces *E. coli* colibactin production, which in turn is inhibitory to *S. aureus in vitro* and *in vivo*, informing the microbial ecology at play during polymicrobial wound infections. Additionally, we report that the BarA-UvrY TCS indirectly regulates *pks* gene expression via the Csr system. The antimicrobial spectrum of colibactin is not limited to *S. aureus* species as previously reported (Fais, Cougnoux et al. 2016), but extends to all the Staphylococal species we tested. However, colibactin bacterial killing appears limited to Staphylococcal genus (Fais, Cougnoux et al. 2016, Keogh, Tay et al. 2016) for reasons we don’t understand at this time, raising the possibility of colibactin-related compounds as a narrow-spectrum anti-Staphylococcal therapeutic. Its genotoxicity toward mammalian cells notwithstanding, colibactin has also been enigmatic to purify at useful yields (Fais, Delmas et al. 2018), complicating its optimization as a therapeutic. Nonetheless, this work underscores the importance of the BarA-UvrY two component system in the regulation of the *pks* island, which could potentially be a therapeutic target to inhibit colibactin synthesis.

## Acknowledgements

This work was supported by the National Research Foundation and Ministry of Education Singapore under its Research Centre of Excellence Programme, by the Ministry of Education Singapore under its tier 2 program (MOE2014-T2-2-124), and by NIH NIAID R21 AI126023-01. We thank Daniela Moses and colleagues for performing library preparation, whole genome sequencing and RNA-Seq; Swaine Chen for providing plasmid pKM208, pSLC-217 and pTrc99a. We thank the NTU Protein Production Platform (www.proteins.sg) for the cloning, expression test, and purification of ClbS protein.

## Author contributions

J.J.W. and K.A.K. designed experiments, analysed data and prepared the manuscript. J.J.W., K.K.L.C., F.K.H., B.C.M.H., and P.Y.C. performed experiments and analysed data. K.K.L.C. analysed RNA-seq data. J.J.W., D.K., and K.A.K. conceptualised the study. R.N. and C.F.L. synthesized *N*-myristoyl-D-Asn for this study. J.C. provided plasmids. All authors reviewed the manuscript.

## STAR methods

### Bacteria strains and growth conditions

All strains and plasmids used in this study are listed in **Table S2**. Both *E. coli* and *S. aureus* were grown in Tryptic Soy Broth (TSB; Merck, Singapore) at 37 °C either with shaking at 200 RPM or under static conditions to late stationary phase. Overnight cultures were normalized to 1-2 × 10^8^ colony forming units (CFU)/ mL by washing the cell pellets twice with phosphate buffered saline (PBS) and then normalized to optical density (OD_600nm_) of 0.4 (*E. coli*) and 0.5 (*S. aureus*) by diluting in PBS.

### Generation of deletion mutants

*E. coli* UTI89 mutants were generated using the positive-negative selection system as described previously (Khetrapal, Mehershahi et al. 2015). Briefly, the first recombination requires amplification of the positive-negative selection cassette (Kan_RelE) from the plasmid pSLC-217 via PCR. Primers contained 50 bp homology sequence that upstream or downstream to the target gene. *E. coli* UTI89 carrying the pKM208 plasmid were induced with 1 mM IPTG and made electro-competent. The competent cells were transformed with 1 μg of PCR product via electroporation. The electroporated cells were recovered in LB at 37 °C for 3 hr with shaking, followed by static incubation for 1 hr. The transformed cells were plated on LB agar plates supplemented with 50 μg/mL kanamycin to select for cells with the Kan_RelE selection cassette inserted into the target gene. The second recombination requires the amplification of 500 bp of DNA sequence that is upstream and downstream of the target gene and stitching them together. For the second recombination, *E. coli* UTI89 with the positive-negative cassette inserted into the target gene and carrying the pKM208 plasmid were induced with 1 mM IPTG and made electro-competent. The electrocompetent cells were transformed with 1 μg of the stitching fragment via electroporation. After recovery, the cells were plated on M9 agar plates supplemented with 0.2 % rhamnose. The resulting knockout mutants were confirmed by colony PCR (see **Table S3** for primers used in this study).

### Generation of plasmids

To create the ClbS expression vector, plasmid pJC-2343 was linearized with primers (InFusion_Vector_F/InFusion_Vector_R) and ClbS was amplified with primers (InFusion_ClbS_F/ InFusion_ClbS_R) using *E. coli* UTI89 genomic DNA as a template. PCR was performed using Q5^®^ High-Fidelity DNA polymerase (New England Biolabs, United States) according to the manufacturer’s protocol. Thereafter, PCR purification was performed using Wizard^®^ SV Gel and PCR Clean-Up System (Promega, United States) following the manufacturer’s protocol. ClbS DNA was inserted into the linearized plasmid using In-Fusion HD Cloning system (Takara, Japan). The infusion product was used to transform into Stellar™ Competent Cells (Takara, Japan). The vector was linearized by inverse PCR with outward directed primers (SodA_RBS_F/ SodA_RBS_R) containing the SodA RBS and re-ligated using Kinase, Ligase, DpnI (KLD) mix (New England Biolabs, United States) (Malone, Boles et al. 2009). The plasmid pJC-2343-ClbS was extracted from Stellar™ competent cells using the Monarch^®^ Plasmid Miniprep Kit and used to transform *S. aureus* USA300 LAC via phage generalized transduction. The sarA P1 promoter was amplified from NCTC 8325 with primers JCO 1141 + JCO 1142 and cloned into pJC1213 at SphI and PstI to generate pJC2343 (Chen, Ram et al. 2015). Primers used are shown in **Table S3**. Successful expression of ClbS in *S. aureus* USA300 LAC was verified by Western blot using guinea pig polyclonal antisera against ClbS and an anti-Guinea pig-HRP secondary antibody for detection.

To create vectors for expression of *csrA* and *csrB*, the respective genes were amplified using *E. coli* UTI89 genomic DNA as a template and primers containing NcoI and HindIII restriction sites. PCR was performed using Q5^®^ High-Fidelity DNA polymerase (New England Biolabs, United States) according to the manufacturer’s protocol, using primers (OEcsrA_F/OEcsrA_R) for the *csrA* insert and primers (OEcsrB_F/ OEcsrB_R) for the *csrB* insert. PCR purification was performed using Wizard^®^ SV Gel and PCR Clean-Up System (Promega, United States) in accordance with the manufacturer’s protocol. Thereafter, the vector pTrc99A and the PCR products were digested with NcoI-HF^®^ and HindIII-HF^®^ (New England Biolabs, United States) according to the manufacturer’s protocol. The vector and insert were ligated with T4 DNA ligase (New England Biolabs, United States) following the manufacturer’s protocol. The ligated product was used to transform Stellar™ Competent Cells (Takara, Japan). The plasmids pTrc99A-CsrA and pTrc99A-CsrB were extracted from Stellar™ competent cells using the Monarch^®^ Plasmid Miniprep Kit and used to transform electrocompetent *E. coli* UTI89. The resulting knockout mutants were confirmed by colony PCR. Primers used in this study are listed below in **Table S3**.

### Generation of polyclonal antisera

Recombinant protein fragments were designed, expressed, and purified using the Protein Production Platform (NTU, Singapore) as previously described (Afonina, Lim et al. 2018). The ClbS target comprised of amino acid residues 2 to 166 from NCBI RefSeq accession no. ABE07674.1 and were cloned into pNIC28-Bsa4 with an N-terminal His tag followed by a TEV protease cleavage site. Polyclonal antisera were generated commercially (SABio, Singapore) by immunization of guinea pigs with purified recombinant ClbS. Specificity of the immune sera was confirmed by the absence of signal on Western blots of whole-cell lysates from wild-type *S. aureus* USA300 LAC with vector control.

### Macrocolony biofilm assay

*E. coli* and *S. aureus* were grown to late stationary phase and normalized as described above. Normalized cultures of *E. coli* and *S. aureus* were mixed at a 1:1 ratio for mixed species macrocolony inocula or diluted twice with PBS for single species macrocolony inocula. 5 μL of each mixture were spotted on TSB supplemented with 1.5 % (w/v) agar. Macrocolonies were grown at 37 °C to the required timepoint. Thereafter, the macrocolonies were harvested using a sterile blade and resuspended in PBS. For enumeration of viable CFU of each strain, the resuspension was plated on medium to select for *E. coli* (McConkey; Merck Singapore) or *S. aureus* (TSB supplemented with colistin and nalidixic acid; 5 μg/mL each).

### Planktonic co-culture assay

*E. coli* and *S. aureus* were grown to late stationary phase and normalized as described above. Normalized cultures of *E. coli* and *S. aureus* were mixed at a 1:1 ratio for mixed cultures or diluted twice with PBS for single cultures. 5 μL of each mixture was inoculated in 5 mL of TSB broth and grown at 37 °C with shaking at 200 RPM or under static conditions. At specific timepoints, 200 μL of the culture was sampled for enumeration of viable CFU before performing serial dilution and plating on selective medium to select for *E. coli* and *S. aureus*.

### RNA extraction from macrocolonies

Single species and mixed species macrocolonies were grown for 6 hours followed by RNA extraction. The macrocolony was first resuspended in TRIzol Reagent (Ambion) and physical cell lysis was performed using Lysing Matrix B (MP Biomedicals). Thereafter, nucleic acids were purified via chloroform extraction followed by isopropanol precipitation. To remove DNA, DNase treatment was performed using the TURBO DNA-free kit (Ambion, USA). Ribosomal RNA (rRNA) was depleted from the samples using the RIBO-Zero Magnetic Bacterial Kit (Epicentre). RNA was converted to cDNA using the NEBNext RNA First Strand Synthesis Module and NEBNext Ultra Directional RNA Second Strand Synthesis Module (New England Biolabs, USA). Library preparation was performed by the SCELSE sequencing facility and sequenced via Illumina Miseq2500 machine as 250 bp paired reads.

### Transcriptomic analysis

RNA sequencing reads were trimmed via BBMap tools (Bushnell, 2016). The trimmed reads were mapped to *S. aureus* HG001 (GenBank assembly accession GCA_000013425.1) and *E. coli* UTI89 reference genome (GenBank assembly accession GCA_000013265.1) using BWA (version 0.7.15-r1140) (Li and Durbin 2009, Nagalakshmi, Waern et al. 2010). Reads were mapped to predicted open reading frames to each reference genome using HTSeq (Anders, Pyl et al. 2015). Gene expression analyses were done in R (version 3.4.4) using Bioconductor package, *edgeR* (Robinson, McCarthy et al. 2010). Gene expression differences were considered significant if the false discovery rate (FDR) was below 0.05. Annotation of genes was done using Kyoto Encyclopedia of Genes and Genomes (KEGG). The raw and processed data for the RNA-seq can be found under the GEO accession number: GSE190571.

### Generation of *E. coli* transposon mutant library

*E. coli* UTI89 were made electrocompetent, achieving a transformation efficiency of 10^7^-10^9^ CFU/μg of DNA. Briefly, pre-warmed SB medium (Tryptone, 30 g/L; yeast extract, 20 g/L; MOPS, 10 g/L) were inoculated with overnight cultures at a 1:250 ratio and incubated at 37 °C with shaking at 200 RPM to mid-log phase (OD_600nm_ 0.8-0.9). The cultures were then chilled on ice for 15 min before washing the cell pellets 3 times in ice cold 10% glycerol. The cell pellets were resuspended in 1 ml of 10% glycerol and aliquoted into 50 μL aliquots. The aliquots were flash frozen in liquid nitrogen and stored at -80 °C. A transposon library of *E. coli* UTI89 was generated with the EZ-Tn5™ <R6Kγori/KAN-2>Tnp Transposome™ Kit (Epicentre®), according to the manufacturer’s protocol. Following transformation, the electroporated cells were allowed to recover at 37 °C for 1 hr in SOC media (Yeast extract, 5 g/L; Tryptone, 20 g/L; 10 mM NaCl; 2.5 mM KCl; 10 mM MgCl_2_; 10 mM MgSO_4_; and 20 mM glucose). Finally, the electroporated cells were diluted in PBS to achieve approximately 100 CFU/plate. The diluted cells were spread on Miller’s LB 1.5% (w/v) agar plates supplemented with 50 μg/mL kanamycin and incubated overnight at 37 °C.

### Transposon library screen

Individual mutants of the *E. coli* UTI89 transposon library were inoculated in 200 μL of LB media in 96-well plates and incubated at 37 °C statically overnight. A *S. aureus* USA300 LAC-GFP overnight culture was normalized as described above and diluted 100-fold to a final volume of 200 μL in 96-well plates before 3 μL of each UTI89 mutant were transferred into each well. Finally, 3 μL of the mixed cultures were spotted onto TSB agar and incubated at 37 °C for 48 hours. A primary screen was conducted based on fluorescence intensity within the macrocolony, indicative of viable GFP-expressing *S. aureus*. Subsequently, mutants from the primary screen were validated by macrocolony biofilm assays and growth kinetic assays before whole genome sequencing was performed to identify the location of the transposon.

### Solid phase synthesis of N-myristoyl-D-Asn synthesis

The synthesis of *N*-myristoyl-D-Asn (NMDA) was performed as previously described (Vizcaino, Engel et al. 2014), with modifications. All the solvents and reagents were purchased from commercial suppliers and used without further purification. N2-Fmoc-N4-trityl-D-asparagine [Fmoc-D-Asn(Trt)-OH], 2-chlorotrityl chloride resin (1.0mmol/g, 100∼200mesh, 1%DVB) and PyBOP were purchased from GL Biochem (Shangai) Ltd. 1H NMR was recorded on a Bruker 400 MHz spectrometer at 298 K. All chemical shifts were quoted in ppm and coupling constants were measured in Hz. Electrospray ionization mass spectrum (ESI-MS) of NMDA was measured in negative mode on a Thermo LTQ XL system.

#### Pre-activation of 2-chlorotrityl chloride resin

A polystyrene resin carrying a 2-chlorotrityl chloride linker (500 mg, 0.75 mmol, 1.5 mmol/g) was placed into a 50 mL polypropylene syringe fitted with a polyethylene porous frit (20 µm). The resin was swollen with dry DMF (3 × 10 mL). After removal of DMF, a solution of thionyl chloride (200 µL, 7.0 µmol) in DMF (5 mL) was added and the reaction mixture stirred for 1 h. The re-activated 2-chlorotrityl chloride resin (**Supplementary Figure 3:1**) was washed with DMF (3 ×10 mL) and dry dichloromethane (DCM, 3×10 mL).

#### Loading of Fmoc-D-Asn(Trt)-OH on 2-chlorotrityl chloride resin

Fmoc-D-Asn(Trt)-OH (**Supplementary Figure 3:2**, 3 equiv.) was mixed with 2-chlorotrityl chloride resin (**Supplementary Figure 3:1**) in anhydrous DCM (10 mL), followed by addition of N,N-diisopropylethylamine (DIPEA, 3 equiv.). The mixture was shaken for 30 min at room temperature. The resin was washed with DMF (10 mL) and the remaining reactive chloride groups were quenched with a solution of DCM:MeOH:DIPEA (5 mL, 80:15:5), followed by washing with DMF (3 × 5 mL) to yield the resin (**Supplementary Figure 3:3**).

#### Fmoc deprotection

To the resin (**Supplementary Figure 3:3**, 0.75mmol) pre-swollen in DCM was added 20% piperidine in DMF (10mL) and the reaction mixture was shaken for 10 min. The solution was drained, and the resin was washed with DMF (x3), DCM (x3). This procedure was repeated twice to obtain the resin (**Supplementary Figure 3:4**). Myristic acid (**Supplementary Figure 3:5**, 3 eqiv.) and PyBOP (6 equiv.) were dissolved in DMF/DCM (50/50). DIPEA (8 eq) was added to the mixture to activate the carboxylic acid. The solution was added to the resin (**Supplementary Figure 3:4**) and the mixture was shaken for 1 h at room temperature. Completion of the coupling reaction was checked using the Ninhydrin test. The solution was drained and the resin was washed with DMF (3 times), DCM (3 times) successively to give the resin (**Supplementary Figure 3:6**).

#### Cleavage of NMDA from the resin

To the resin (**Supplementary Figure 3:6**) was added the cleavage mixture TFA/H2O/TIS (95%/2.5%/2.5%, 5mL) and the mixture was shaken for 3 h at room temperature. The resin was removed by filtration and the resin was washed with the cleavage mixture once (2.5 mL). To the combined filtrate was added dropwise cold diethyl ether to precipitate the crude NMDA. The precipitate was collected after centrifugation and the diethyl ether decanted. This solid was washed with cold diethyl ether three times (20-30 mL x3) using the centrifugation procedure. The crude product was purified by semi-preparative reverse-phase-HPLC. Semi-preparative RP-HPLC was preformed using a Shimadzu HPLC system equipped with a Phenomenex jupiter-C18 RP column (10 × 250 mm, 5 μm) with a flow rate of 2.5 mL per minute, eluting using a gradient of buffer B (90% acetonitrile, 10% H2O, 0.045% TFA) in buffer A (H2O, 0.045% TFA). The combined pure NMDA fractions after HPLC purification were lyophilized to afford N-myristoyl-D-asparagine in powder form.

#### Compound characterization

The obtained pure compound was characterized by 1H NMR (**Supplementary Figure 4**). (400 MHz DMSO-d6): 12.46 (br,1H COOH), 7.96 (d, J = 8 Hz,1H, C(O)NHCH), 7.32 (s, 1H, C(O)NH2), 6.87 (s, 1H, C(O)NH2), 4.47-4.51 (m, 1H,NHCH), 2.52 (dd, J = 5.7, 15.5 Hz, 1H, CH2C(O)NH2), 2.41 (dd, J = 7.2, 15.5 Hz, 1H,CH2C(O)NH2), 2.07 (t, J = 7.3 Hz, 2H, C(O)CH2CH2),1.48-1.42 (m, 2H, C(O)CH2CH2), 1.28-1.19 (m, 20H, myristoyl-CH2), 0.85 (t, J = 6.8 Hz, 3H, CH2CH3). ESI-MS: m/z [M–H]?calculated for C18H33N2O4? 341.24 (isotopic), observed 341.39 (**Supplementary Figure 5**).

### N-myristoyl-D-Asn growth inhibition assay

Overnight cultures of *S. aureus* were normalized to OD_600nm_ of 0.4 and diluted 100-fold. Thereafter, 8 μL of the diluted cultures were inoculated into 96-well plates containing 200 μL of TSB media supplemented with NMDA at 100 μM, 300 μM and 600 μM. For the vehicle control, DMSO was supplemented to 1% (v/v). The plates were incubated at 37 °C in a Tecan Infinite© M200 Pro spectrophotometer. Absorbance readings at 600 nm were taken every 15 min for 12 hours.

### Cell permeability assay

Cell permeability was determined using propidium iodide (PI) staining and flow cytometry. *S. aureus* USA300 LAC overnight cultures were sub-cultured into fresh TSB media and allowed to grow to mid-log phase at OD_600nm_ of 0.6. Subsequently, the cultures were treated with DMSO (1%), 100 μM palmitoleic acid and 600 μM NMDA and incubated at 37 °C for 15 min. After the treatment, the cells were washed twice with PBS and stained with PI Buffer (Abcam) for 30 min at room temperature. Flow cytometry was performed with flow cytometer Fortessa X to identify the population of *S. aureus* stained positive for PI after treatment with compounds. Data analysis was performed using FlowJo, version 10.

### RNA extraction and real time quantitative PCR (RT-qPCR)

Macrocolonies were grown as described above. RNA from macrocolonies was extracted using the RNeasy^®^ Mini Kit (Qiagen, United States) according to the manufacturer’s protocol. Genomic DNA was removed by DNase treatment (TURBO DNA-free Kit, Ambion). RNA and DNA were quantified using Qubit™ RNA Assay Kit and Qubit™ dsDNA HS Assay kits (Invitrogen, United States). RNA quality was analyzed using Agilent RNA ScreenTape (Agilent Technologies, United States). RNA samples with minimum RIN value of 7.5 and DNA contamination of not more than 10% were converted to cDNA using SuperScript™ III First-strand Synthesis Supermix (Invitrogen, United States) with accordance to the manufacturer’s protocol. RT-qPCR reaction mix was prepared using KAPA SYBR^®^ FAST qPCR Kit Master Mix (2X) Universal (Kapa biosystem, United States) and ran on a StepOnePlus™ Real-Time PCR System (Applied Biosystems, USA). Primers GyrA_F/GyrA_R were used to amplify *gyrA* (Housekeeping gene), primers ClbA_F/ClbA_R were used to amplify *clbA*, primers ClbB_F/ClbB_R were used to amplify *clbB*. Primers are found in **Table S4**.

### Mouse model of polymicrobial wound infection

Bacteria was grown as described above, normalized to 10^6^CFU/10 µl, and used to infect wounds of C57BL/6 mice (Male, 7-8 weeks old; InVivos, Singapore) as previously described (Chong, Tay et al. 2017). Briefly, the animals were anesthetized with 3% isoflurane. Dorsal hair was shaven and fine hair was removed after the application of Nair™ cream (Church and Dwight Co, Charles Ewing Boulevard, USA) and shaved using a scalpel blade. The skin was disinfected with 70 % ethanol and a full-thickness wound was created with a 6 mm biopsy punch (Integra Miltex, New York, USA). The wounds were inoculated with 10 μL of the respective inoculum (*E. coli*, 1-2 × 10^6^ CFU; *S. aureus* 1-2 × 10^6^ CFU; Mixed 1-2 × 10^6^ CFU each). Thereafter, the wound site was sealed with a transparent dressing (Tegaderm™ 3M, St Paul Minnesota, USA). At the indicated timepoints, mice were euthanized, and the wounds were excised and homogenized in 1 mL PBS. Viable bacteria in the wound homogenates were enumerated by plating onto selective media for *E. coli* (McConkey; Merck Singapore) and *S. aureus* (MRSA*Select*™ II Agar; Biorad USA). All animal experiments were performed with approval from the Institutional Animal Care and Use Committee (IACUC) in Nanyang Technological University, School of Biological Sciences under protocol ARF-SBS/NIE-A19061.

