## Supplementary Figures for "*Escherichia coli* BarA-UvrY regulates the *pks* island and kills Staphylococci via the genotoxin colibactin"

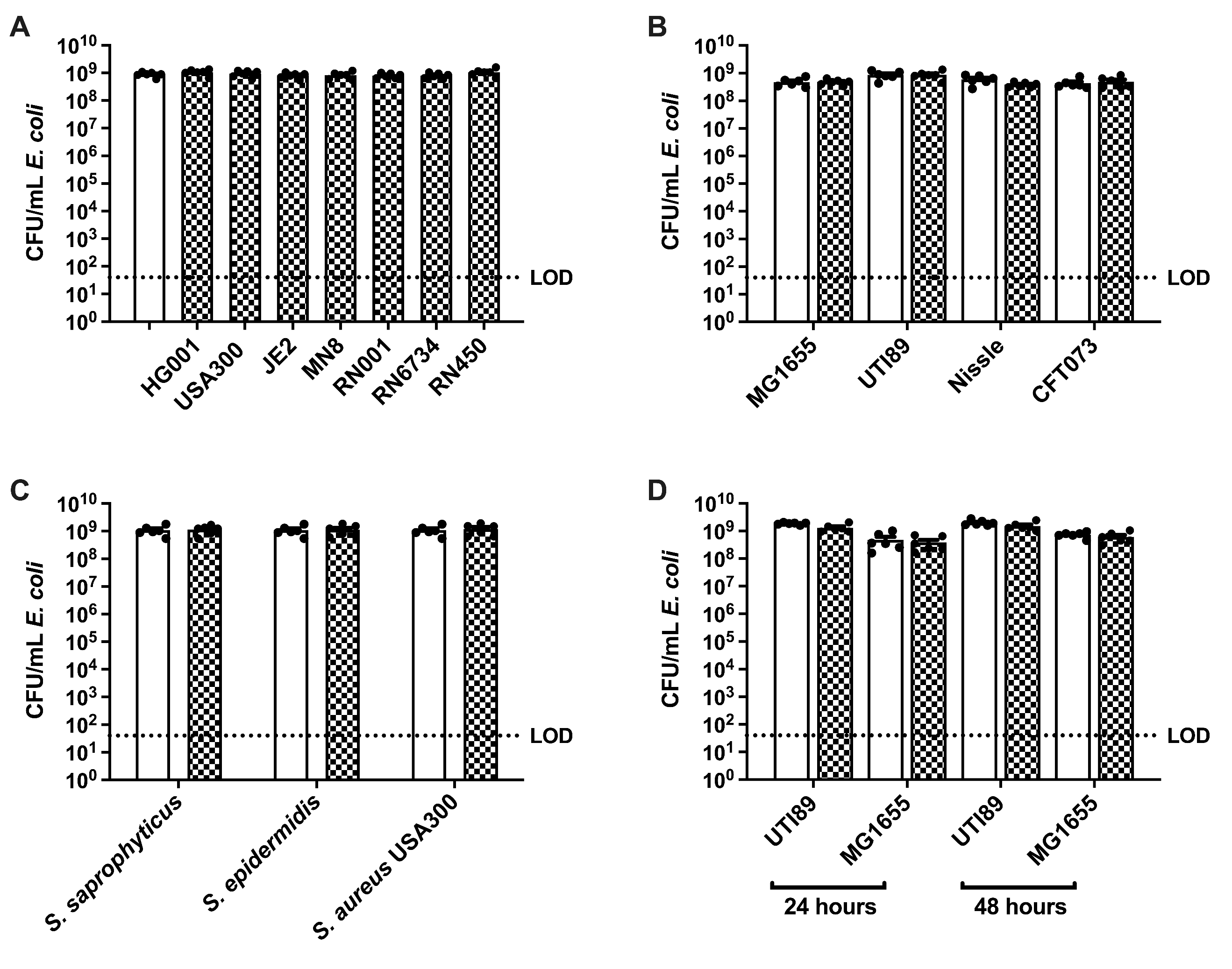

**Supplementary Figure 1. *E. coli* CFU are unchanged upon growth with *S. aureus*. (A)** Enumeration of *E. coli* UTI89 grown alone or together with different strains of *S. aureus* in macrocolonies for 24 h. N=6 independent experiments. **(B)** Enumeration of indicated strains of *E. coli* from single species or mixed species macrocolonies containing *S. aureus*, at 24 hours. N=6 independent biological experiments. **(C)** Enumeration of *E. coli* UTI89 from single species or mixed species macrocolonies co-cultured with indicated Staphylococcal species for 24 hours. N=6 independent experiments. **(D)** Enumeration of *E. coli* strains UTI89 or MG1655 after planktonic growth alone or mixed with *S. aureus* for 24 or 48 hours. N=6 independent experiments (**A-D**) Data from single species macrocolonies or planktonic cultures are indicated with open bars, and data from mixed species (all inoculated at a ratio of 1_EC_:1_SA_) macrocolonies are indicated with checked bars. Individual data points from each biological replicate are indicated with closed circles. No statistical significance was detected for any of the comparisons shown. Error bars represent SE from the mean.

| Table S1. List of Transposon Mutants Identified from Transposon Library Screen | | | | |
| --- | --- | --- | --- | --- |
| Gene Name | Gene Locus | Description | | # of mutants |
| *clbA* | UTI89_RS10835 | Colibactin biosynthesis phosphopantetheinyl transferase ClbA | 2 | |
| *clbR* | UTI89_RS10830 | Colibactin biosynthesis LuxR family transcriptional regulator ClbR | 3 | |
| *clbB* | UTI89_RS10825 | Colibactin hybrid non-ribosomal peptide synthetase/type I polyketide synthase ClbB | 24 | |
| *clbD* | UTI89_RS10815 | Colibactin biosynthesis dehydrogenase ClbD | 12 | |
| *clbE* | UTI89_RS10810 | Colibactin biosynthesis aminomalonyl-acyl carrier protein ClbE | 5 | |
| *clbF* | UTI89_RS10805 | Colibactin biosynthesis dehydrogenase ClbF | 6 | |
| *clbG* | UTI89_RS10800 | Colibactin biosynthesis acyltransferase ClbG | 3 | |
| *clbH* | UTI89_RS10795 | Colibactin non-ribosomal peptide synthetase ClbH | 4 | |
| *clbI* | UTI89_RS10790 | Colibactin polyketide synthase ClbI | 5 | |
| *clbJ* | UTI89_RS10785 | Colibactin non-ribosomal peptide synthetase ClbJ | 8 | |
| *clbK* | UTI89_RS10780 | Colibactin hybrid non-ribosomal peptide synthetase/type I polyketide synthase ClbK | 8 | |
| *clbN* | UTI89_RS10765 | Colibactin non-ribosomal peptide synthetase ClbN | 5 | |
| *clbO* | UTI89_RS10760 | Colibactin polyketide synthase ClbO | 1 | |
| *clbP* | UTI89_RS10755 | Pre-colibactin peptidase ClbP | 4 | |
| *clbQ* | UTI89_RS10750 | Colibactin biosynthesis thioesterase ClbQ | 4 | |
| *uvrY* | UTI89_RS10285 | Two-component system response regulator UvrY | 1 | |
| *barA* | UTI89_RS15370 | Two-component sensor histidine kinase BarA | 4 | |
| *cysP* | UTI89_RS13485 | Thiosulfate binding protein | 1 | |
| *aroH* | UTI89_RS09215 | Phospho-2-dehydro-3-deoxyheptonate aldolase | 1 | |
| *dnaJ* | UTI89_RS00080 | Chaperone with DnaK, heat shock protein | 1 | |
| *waaL* | UTI89_RS20145 | Putative lipid A-core surface polymer ligase | 2 | |
| *htpG* | UTI89_RS02395 | Chaperone Hsp90, heat shock protein | 2 | |
| *wbdM* | UTI89_RS11255 | Putative glycosyltransferase | 1 | |
| *rfaQ* | UTI89_RS20185 | Lipopolysaccharide core biosynthesis glycosyl | 1 | |

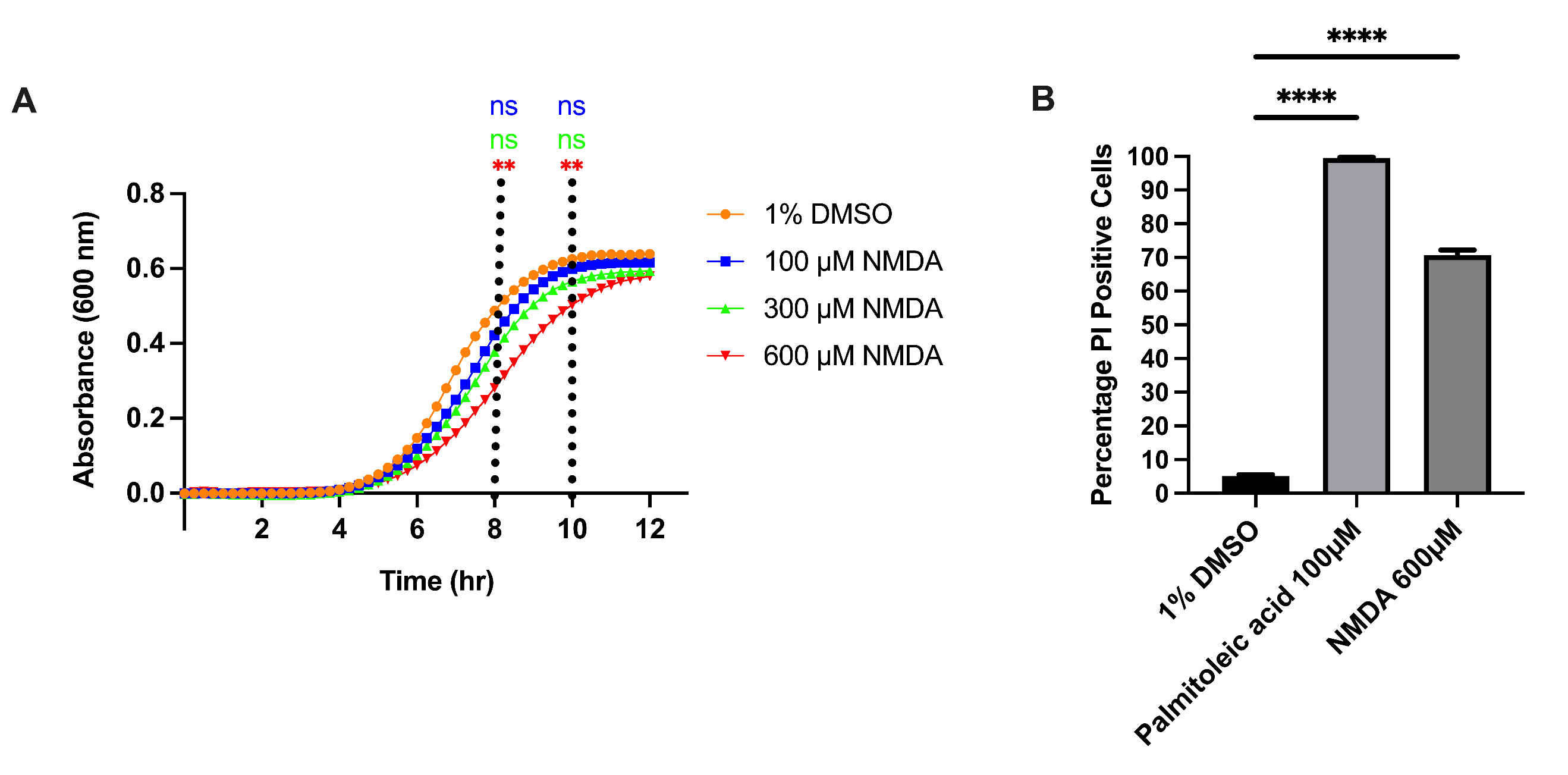

**Supplementary Figure 2. *N*-myristoyl-D-Asn compromises *S. aureus* membrane integrity. (A)** Growth curves of WT *S. aureus* USA300 LAC grown in TSB supplemented with 100 μM, 300 μM and 600 μM of NMDA. An equal concentration of DMSO (1%) was used as the vehicle control. Each data point represents the mean measurement from 3 biological replicates, each the average of 4 technical replicates. Statistical analysis was done using Kruskall-Wallis Test with Dunn’s post-test to correct for multiple comparisons. **p< 0.01. **(B)** Percentage of PI positive *S. aureus* cells after treatment with 1% DMSO, 100 μM palmitoleic acid (16:9) or 600 μM NMDA. Statistical significance was determined by One-way ANOVA with Dunnett’s test for multiple comparison. ****p< 0.0001. Error bars represent SE from the mean.

| Table S2. List of bacterial strains and plasmids used in this study | | |
| --- | --- | --- |
| Bacterial Strains | **Description** | **References** |
| *Escherichia coli* | | |
| UTI89 | Uropathogenic clinical isolate | (Chen et al. 2006) |
| CFT073 | Uropathogenic clinical isolate | (Welch et al. 2002) |
| MG1655 | *E. coli* K-12 strain, LPS mutant | (Blattner et al. 1997) |
| Nissle 1917 | Non-pathogenic gut isolate | (Scaldaferri et al. 2016) |
| UTI89 ∆*pks* | *pks* island knockout mutant | This study |
| UTI89 ∆*barA* | *barA* knockout mutant | This study |
| UTI89 ∆*uvrY* | *uvrY* knockout mutant | This study |
| UTI89-pCsrA | UTI89 (pTrc99a-CsrA) | This study |
| UTI89-pCsrB | UTI89(pTrc99a-CsrB) | This study |
| UTI89 ClbR^mut^ | UTI89 strain with GGA bases mutated in *clbR* gene | This study |
| *Staphylococcus aureus* | | |
| RN001 | *S. aureus* *rsbU* mutant | (Novick 1967) |
| HG001 | RN001 derivative, *rsbU* repaired | (Herbert et al. 2010) |
| RN6734 | 8325-4 derivative, *agr*^+^ | (Vojtov, Ross, and Novick 2002) |
| MN8 | Clinical isolate of toxic shock syndrome | (Schlievert and Blomster 1983) |
| USA300 LAC | Community-associated MRSA USA300 | (Miller et al. 2005) |
| JE2 | USA300 LAC, p01 and p03 cured | (Fey et al. 2013) |
| RN450 | Prophage cured *S. aureus* strain 8325 | (Novick 1967) |
| USA300 LAC-GFP | USA300 LAC, pALC1420 GFP+ | (Pang et al. 2010) |
| RN4220 | *S. aureus* strain NCTC 8325-4, *sau1*^-^ *hsdR*^-^, restriction deficient | (Kreiswirth et al. 1983) |
| USA300-ClbS | USA300 LAC (pJC-2343-ClbS) | This study |
| Plasmids | | |
| pKM208 | Red recombinase expressing plasmid; Amp^R^ | (Murphy and Campellone 2003) |
| pSLC-217 | P*_rhaB_ relE* cassette; Neo^R^ | (Khetrapal et al. 2015) |
| pTrc99a | *E. coli* cloning vector containing IPTC inducible promoter P*trc*; Amp^R^ | (Amann, Ochs, and Abel 1988) |
| pJC1213 | *S. aureus* vector pT181 replicon, CmR | (Chen et al. 2015) |
| pJC2343 | pJC1213, P*sarA* P1 promoter, CmR | This study |
| pTrc99a-CsrA | *E. coli* cloning vector pTrc99a::*csrA* (NcoI/HindIII); Amp^R^ | This study |
| pTrc99a-CsrB | *E. coli* cloning vector pTrc99a::*csrB* (NcoI/HindIII); Amp^R^ | This study |
| pJC-2343 | *S. aureus* cloning vector containing P*sarA* P1 promoter; Cm^R^ | This study |
| pJC-2343-ClbS | *S. aureus* cloning vector pJC-2343::*clbS*; Cm^R^ | This study |

| **Table S3. List of primers** | | | |
| --- | --- | --- | --- |
| **Primer** | **Target** | **Sequence 5’-3’** | **Purpose** |
| PKS_217_F | *pks* island | ATCGCTGCCCGGAAAAATCATCAGTGGGGAGGCAAACGGTAAGCACCCCGgcttgcagtgggcttacatg | Amplification of the Kan_RelE selection cassette for insertion into the *pks* island |
| PKS_217_R |  | AAAATCAATATTATCGACGGCTCAGAAGTGTCTAGATTATCCGTGGCGATcccatccagtgcaaagctagc |  |
| BarA_217_F | *barA* | GACTTTCTCAATTTAACAGTGTGACCTTAATTGTCCCATAACGGAACTCCgcttgcagtgggcttacatg | Amplification of the Kan_RelE selection cassette for insertion into the BarA |
| BarA_217_R |  | ATCTGAAACCAGCGTCATAAAAATCCGGTTGCTACTCGACAAGACGTCCAcccatccagtgcaaagctagc |  |
| UvrY_217_F | *uvrY* | TGGCTGGCTGGTTACGGTTTTTAAAAACGCTTTTGCGTCAAACTGATCACgcttgcagtgggcttacatg | Amplification of the Kan_RelE selection cassette for insertion into the UvrY |
| UvrY_217_R |  | ACGAATGACTAACTATCAGTAGCGTTATCCCTATTTCTGGAGATATTCCT cccatccagtgcaaagctagc |  |
| PKS_F1 | 500 bp up-stream of the *pks* island | CATGGTTTCCGGGCAATGG | Amplification of stitching fragment 1 |
| PKS_R1 |  | GTGTCTAGATTATCCGTGGCGATCGGGGTGCTTACCGTTTG |  |
| PKS_F2 | 500 bp down-stream of the *pks* island | ATCGCCACGGATAATCTAGACAC | Amplification of stitching fragment 2 |
| PKS_R2 |  | CAGACCAGCTCAGGTTCAG |  |
| BarA_F1 | 500 bp up-stream of *barA* | GCTTTGCGAAACCCCATCTG | Amplification of stitching fragment 1 |
| BarA_R1 |  | GCTACTCGACAAGACGTCCAGGAGTTCCGTTATGGGACAATTAAGG |  |
| BarA_F2 | 500 bp down-stream of *barA* | TGGACGTCTTGTCGAGTAGC | Amplification of stitching fragment 2 |
| BarA_R2 |  | GCCGCACCGGGTAAAATC |  |
| UvrY_F1 | 500 bp up-stream of *uvrY* | CAGCGCCCTATCTGATATTGC | Amplification of stitching fragment 1 |
| UvrY_R1 |  | CGTTATCCCTATTTCTGGAGATATTCCTGTGATCAGTTTGACGCAAAAGC |  |
| UvrY_F2 | 500 bp down-stream of *uvrY* | AGGAATATCTCCAGAAATAGGGATAACG | Amplification of stitching fragment 2 |
| UvrY_R2 |  | TGCCAGTTCTTGATTATTCTCGG |  |
| InFusion_ClbS_F | *clbS* | CAGGTCGACTCTAGAATGGCTGTTCCATCATCAAA | Amplification of *clbS* for insertion into the *S. aureus* cloning vector pJC-2343 |
| InFusion_ClbS_R |  | GAGCTCGGTACCCGGCTATTCTGCAAGACATTTCTGCA |  |
| InFusion_Vector_F | pJC-2343 | CCGGGTACCGAGCT | Linearization of the *S. aureus* cloning vector pJC-2343 |
| InFusion_Vector_R |  | TCTAGAGTCGACCTGCAGG |  |
| SodA_RBS_F | pJC-2343-ClbS | TGATTATTTATGGCTGTTCCATCATCAAAAG | Linearization of the *S. aureus* cloning vector pJC-2343-ClbS to include the SodA RBS |
| SodA_RBS_R |  | TCCTCCTAAAATTCGAGCTCGGTACCC |  |
| JCO 1141 | *sarA* P1 promoter | CGGGCATGCGCTGATATTTTTGACTAAACCAAATGC | Amplification of the *S. aureus* NCTC 8325 *sarA* P1 promoter to create pJC2343 |
| JCO 1142 |  | TTCCTGCAGGATGCATCTTGCTCGATACATTTGC |  |
| OEcsrA_F | *csrA* | CGAGCACCATGGAAGAAGGAGATATACATATGCTGATTCTGACTCGTCGAG | Amplification of *csrA* for insertion into the *E. coli* cloning vector pTrc99a |
| OEcsrA_R |  | CGAGCAAAGCTTAGTAACTGGACTGCTGGGATTTTTC |  |
| OEcsrB_F | *csrB* | CGAGCACCATGGGAGTCAGACAACGAAGTGAAC | Amplification of *csrB* for insertion into the *E. coli* cloning vector pTrc99a |
| OEcsrB_R |  | CGAGCAAAGCTTAAATAAAAAAAGGGAGCACTGTATTCA |  |

| **Table S4. List of primers used in RT-qPCR** | | | |
| --- | --- | --- | --- |
| **Primer** | **Target** | **Gene Type** | **Sequence 5’-3’** |
| GyrA_F | *gyrA* | Housekeeping | GCTTAACAACCTCTACTCCC |
| GyrA_R | *gyrA* | Housekeeping | CTTCAAGGATATGAGCACGA |
| ClbA_F | *clbA* | Target | TACGTGCAAATATGGCAAAC |
| ClbA_R | *clbA* | Target | CTAATAGCAACGGCTACTGT |
| ClbB_F | *clbB* | Target | TCAAGGTAGCCAATATCACG |
| ClbB_R | *clbB* | Target | ATCCTCATCGTCAAACAACA |

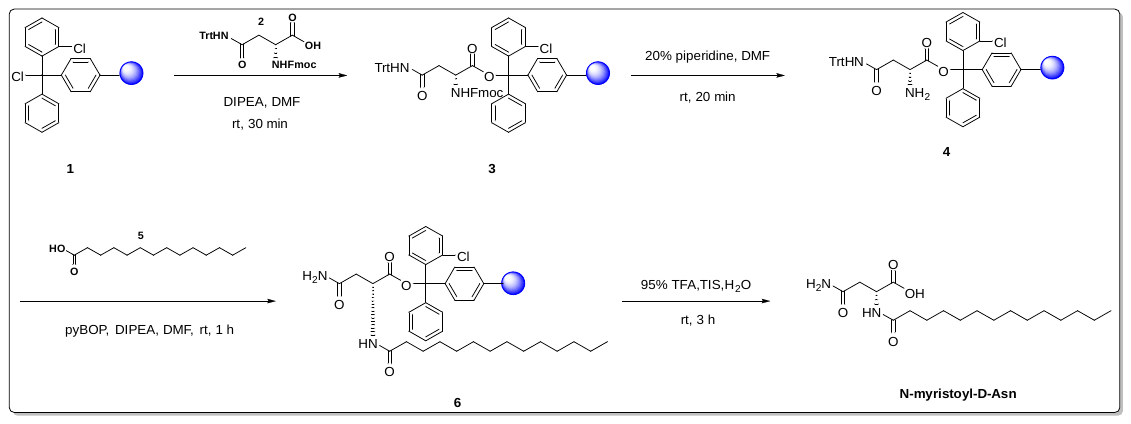

**Supplementary Figure 3. NMDA synthesis scheme.**

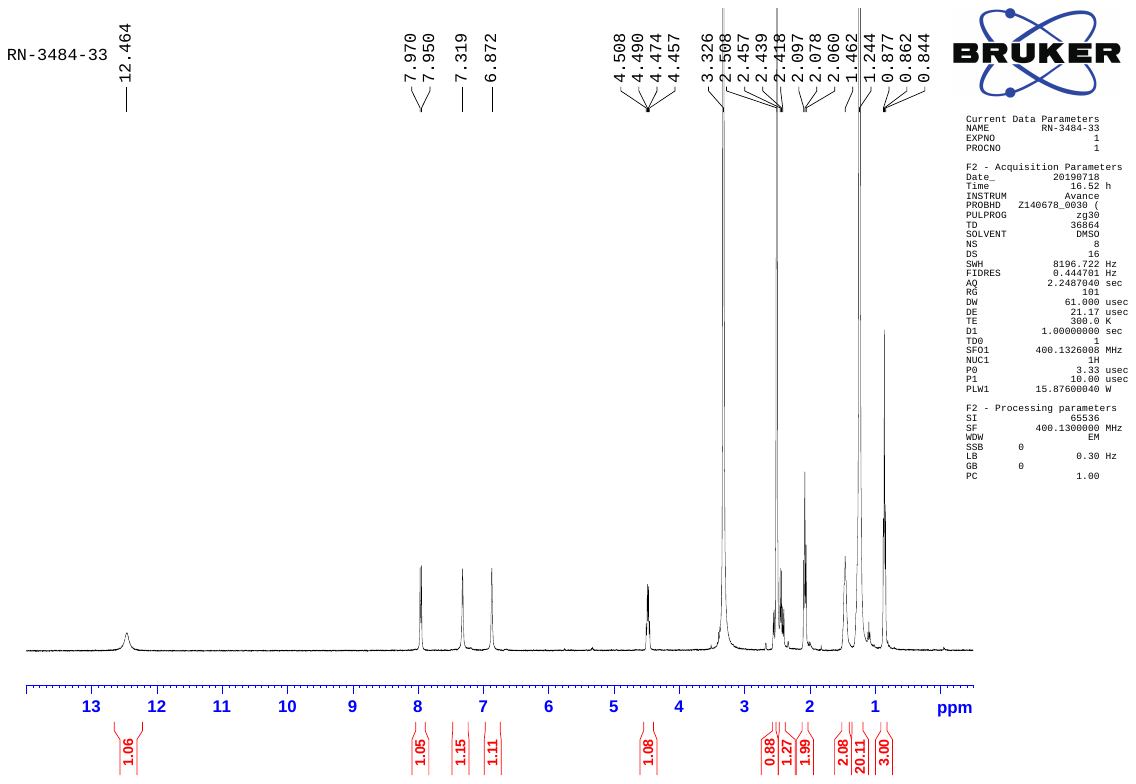

**Supplementary Figure 4. ^1^H-NMR spectrum of N-myristoyl-D-Asn (recorded in DMSo-*d6* at 400 MHz).**

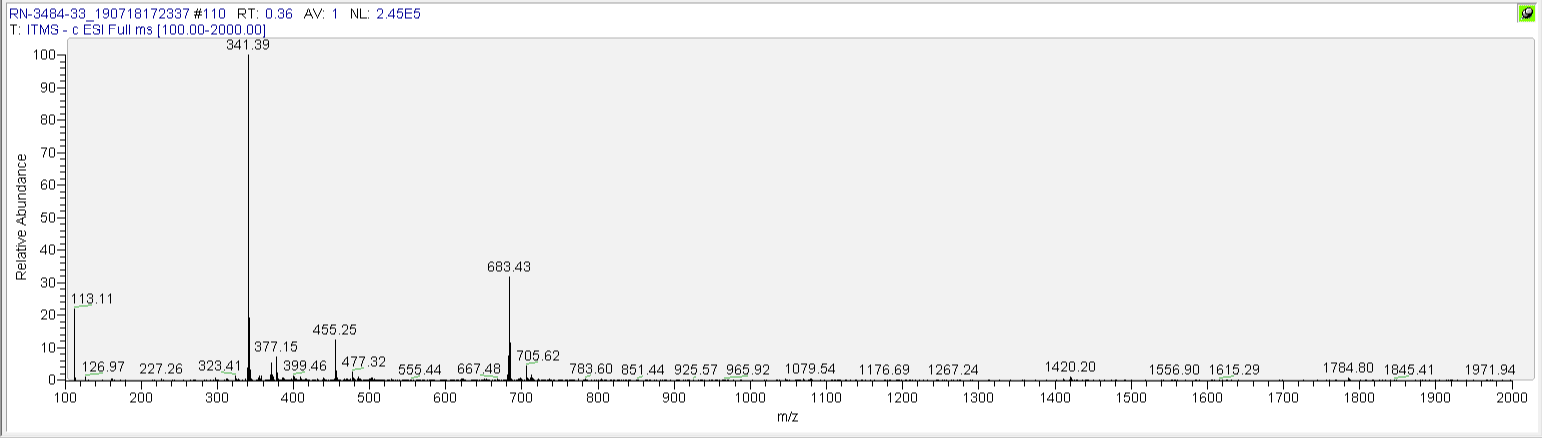

**Supplementary Figure 5. ESI spectrum of N-myristoyl-D-Asn (recorded in negative mode).**
